# Mechanical compression induces persistent bacterial growth during bacteriophage predation

**DOI:** 10.1101/2022.08.12.503793

**Authors:** Guy Mason, Enrique R. Rojas

## Abstract

Although the relationship between bacteria and lytic bacteriophage is fundamentally antagonistic, these microbes not only coexist but thrive side-by-side in myriad ecological environments. The mechanisms by which coexistence is achieved, however, are not fully understood. By examining *Escherichia coli* and bacteriophage T7 population dynamics at the single-cell and single-virion level using a novel microfluidics-based assay, we observed bacteria growing “persistently” when perfused with high-titer bacteriophage. Persistence occurred at a frequency five orders of magnitude higher than is expected from natural selection of bacteriophage-resistant mutants. Rather, the frequency of persistence was correlated with the degree to which the bacteria were mechanically compressed by the microfluidic perfusion chamber. Using a combination of mutagenesis and fluorescent imaging techniques, we found that compression induces persistence by activating the *Rcs* phosphorelay pathway, which results in the synthesis of extracellular capsule that sterically blocks bacteriophage adsorption. Other forms of mechanical stimulation also promoted Rcs activity and persistence. These findings have important implications for our understanding of microbial ecology in many important environments, including the gut and the soil, where bacteria grow in confinement.

**Significance Statement:** Bacteria and bacteriophage form one of the most fundamental and important predator-prey relationships on earth, yet the factors that promote long-term stability of their populations are unknown. Here, we demonstrate that *Escherichia coli* is able to rapidly grow during bacteriophage predation if they are doing so in spatially confined environments. This discovery revises our understanding of bacteria-bacteriophage population dynamics in many real-world environments where bacteria grow in such environments, such as the gut and the soil. Additionally, this result has critical implications for the potential of antibacterial therapies to function during pathogenesis, when bacteria are also mechanically stimulated.

## Introduction

Lytic bacteriophages shape bacterial populations in many important environments, including the gut^1^, the soil^2^, and the ocean^3^. These interactions, moreover, are ancient: bacteriophages and bacteria have co-evolved for more than 3 billion years^4,5^. Thus, despite their antagonistic relationship, the molecular mechanisms that govern the population dynamics of lytic bacteriophage and bacteria have evolved to stabilize their co-existence in the environment.

How is co-existence mediated? One standard view is centered on spatiotemporal evolutionary dynamics^6^. For example, lytic bacteriophage can select for resistant bacterial mutants, which can subsequently lose resistance in bacteriophage-free environments, thereby maintaining populations of susceptible and resistant bacteria across environments^7,8^. This mechanism is essentially an instance of classic Lotka-Volterra predator-prey dynamics^9^. Genetic resistance may rely on mutation of the bacteriophage receptor^10^, constitutive overproduction of extracellular polysaccharide^6^ (which blocks the bacteriophage from its receptor), horizontal gene transfer of restriction modification systems^11^, or acquisition and loss of CRISPR spacers^12^. Alternatively, spatiotemporal population dynamics in the absence of evolution could, in principle, lead to coexistence. For example, theory has demonstrated that bacterial growth on solid media could alone provide a mechanism of co-existence in the absence of genetic resistance if rapidly growing cells on the periphery of a bacterial microcolony shield interior cells^13^.

Environmental factors also influence bacteria-bacteriophage dynamics. In particular, bacteria can survive in the presence of bacteriophage when they adopt a slow-growing, sessile lifestyle within biofilms, where the extracellular matrix slows diffusion of the bacteriophage^14^ and sterically blocks bacteriophage adsorption to the cells^15^. Biofilms may also promote the type of evolutionary dynamics, described above, that lead to bacteria-bacteriophage co-existence^16^. Biofilm synthesis can be induced by a variety of environmental stimuli (starvation, quorum sensing, etc.), although not by bacteriophage itself. In other words, biofilms are an effective but not a specific defense against bacteriophage.

Along these lines, a genetic screen for bacteriophage resistance demonstrated that activation of the Rcs phosphorelay pathway, which mediates a well understood stress-response that induces cells to coat their surface with colanic acid (“capsule”), could in principle provide cross-protection against bacteriophage in response to cell-envelope stress^17^. Capsule synthesis is important for bacterial competitive success in other contexts, for example by preventing colonization of competitors^18^ or blocking Type IV secretion system attack^19^.

There are several key questions that remain unanswered with respect to bacteria-bacteriophage co-existence. First, each of the mechanisms of bacterial survival of bacteriophage predation described above requires either bacteriophage resistance or a reduction in metabolic activity on the part of the bacterium. In either case, it is unclear how bacteriophages avoid extinction since they have relatively short lifetimes in the environment if they are not actively proliferating^20^. Second, lytic bacteriophage and rapidly growing bacteriophage-susceptible bacteria are often found in abundance in the same niche. For example, one lytic bacteriophage active against *Vibrio cholerae* was found at high-titer in 100% of cholera patient stool samples while the load of susceptible bacteria was also very high^21^. This outcome is not consistent with existing models of evolutionary predator-prey dynamics unless the rates of evolution are unrealistically high^22^. Third, the presence of lytic bacteriophage can lead to a counterintuitive increase in the load of *V. cholerae* in a mouse model of cholera^23^. Collectively, these observations point to unknown ecological interactions between the two microorganisms that allow them both to proliferate in close confines.

Here, we demonstrate that growth in confined, two-dimensional environments allows bacteria to grow “persistently” in the presence of a high-titer lytic bacteriophage. We discovered that persistence occurs because physical confinement causes activation of the Rcs phosphorelay pathway, causing capsule synthesis, which reduces the rate of bacteriophage adsorption. Reduced adsorption, in turn, reduces the rate of bacterial lysis and allows bacterial proliferation to balance cell lysis such that susceptible bacteria within bacteriophage-infected microcolonies can grow rapidly and indefinitely. Bacteriophage persistence is a novel innate mechanism by which bacteria can cope with lytic bacteriophage and has several implications for our understanding of bacteriophage-bacteria ecology in real-world environments.

## Results

### Bacteria in confinement grow persistently during bacteriophage predation

Conventionally, bacteria-bacteriophage population dynamics have been measured using bulk-culture assays^24^. Bacteriophage predation of bacterial communities has been observed at the single-cell and -virion level^25^, but not with high temporal and spatial resolution simultaneously. Therefore, in order to examine the physiological basis for bacteria and bacteriophage co-existence, we used microfluidics and single-cell time-lapse microscopy to monitor the spatiotemporal dynamics of bacterial growth under bacteriophage predation for tens of hours continuously (**Fig. 1A,B**).

**Figure 1.**
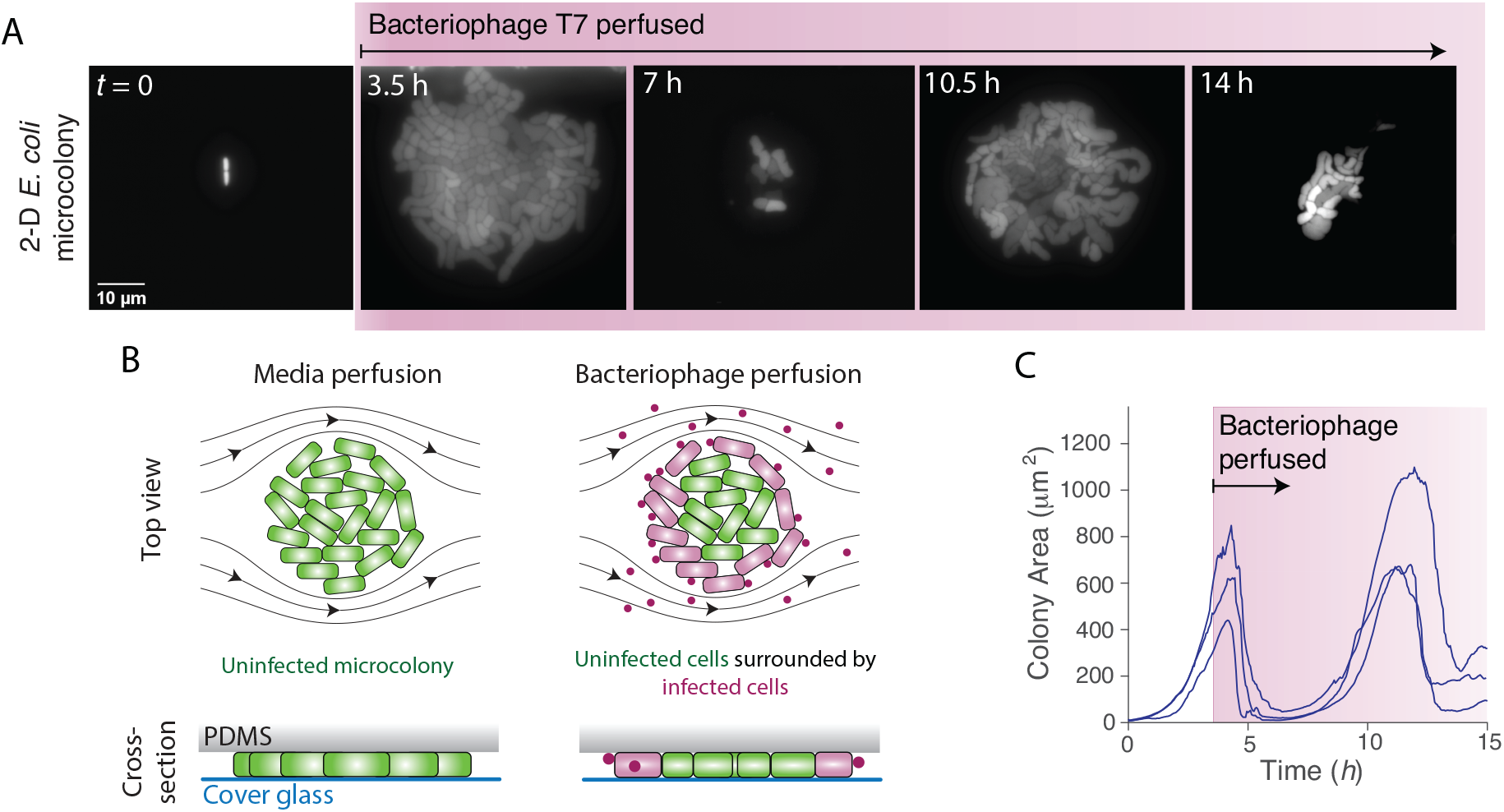
*E. coli* under confinement grows persistently during perfusion of bacteriophage T7. A) Montage of a two-dimensional (2-D) *E. coli* microcolony during perfusion of bacteriophage T7. B) Schematic of the experimental design. C) Colony area versus time for three *E. coli* microcolonies that grew under LB perfusion for 4 hours before being perfused with LB containing bacteriophage T7 indefinitely. Similar results were obtained for 15 experiments.

We first quantified the lysis dynamics of *Escherichia coli* cells by bacteriophage T7. The wild-type bacterial strain we used has no mechanism of bacteriophage immunity. We allowed single bacterial cells to proliferate exponentially for defined periods of time during constant perfusion with rich media (lysogeny broth, LB) before perfusing them indefinitely with media containing high titers of bacteriophage (10^*7*^ ml^-1^). Because within the microfluidic perfusion chamber the cells are trapped between a cover glass and a layer of polydimethylsiloxane (PDMS), single cells grew into two-dimensional microcolonies during this initial phase of growth (**Fig. 1A,B**). The subsequent perfusion of bacteriophage caused colonies to rapidly decrease in size (**Fig. 1C**). Cell lysis began at the colony periphery and progressed toward the center (**Movie S1**), suggesting that peripheral bacterial cells were temporarily protecting central cells by binding and sequestering the bacteriophage (**Fig. 1C**).

Surprisingly, many microcolonies re-established exponential growth after the initial period of rapid lysis despite being continuously perfused with bacteriophage (**Fig. 1A,C**). This “persistent” growth occurred at a frequency (8.6 ± 8.3%; mean ± 1 s.d., *n*=10 experiments) ≈10^5^ higher than would be expected from natural selection of resistant mutants^26,10^. Indeed, these microcolonies were still susceptible to the bacteriophage since they underwent repeated periods of lysis and growth (**Fig. 1B,C,S1A**). Colony growth-rate during the periods of growth was comparable to that of uninfected colonies (**Fig. 1C**), indicating that the bacteria were temporarily immune to the bacteriophage during these periods. Interestingly, the temporal dynamics of microcolony size often exhibited an oscillatory-like pattern that was synchronized between spatially separated microcolonies (**Fig. 1C**), and cells within persistent microcolonies sometime displayed amorphous morphology (**Fig. 1A, Movie S1**), suggesting these dynamics were due to a stereotypical physiological stress response by the bacterium. Persistence occurred after as little as 5 minutes of growth in the perfusion chamber (**Fig. S1A**), and we observed persistent growth for up to 15 hours, which was long as we could continually image the bacteria.

We confirmed that in bulk liquid culture, bacteriophage T7 rapidly kills our wild-type *E. coli* at a rate consistent with classic experiments^10^ (<10^−8^ resistant cells). We therefore hypothesized that persistence specifically depended on confined growth. The rectangular microfluidic perfusion chambers we used are 1500 × 100 µm (*x*-*y*, **Fig. 2A**), and are engineered with a height gradient (0.5 ≲ *h* ≲ 1.5) along their long axis (*x*-axis, **Fig. 2B**), which allows bacterial cells to become selectively trapped at positions where their diameter matches the chamber height. However, with lower probability some cells become trapped in areas of the chamber with heights less than their diameter (**Fig. 2A,C**). The frequency at which this occurs depends on the flow rate used to load the cells into the chamber, allowing us to deliberately wedge cells into very narrow areas of the chamber. Interestingly, although cell density increased with chamber height, the probability for a cell to become persistent strongly decreased with height (**Fig. 2C,D, Movie S2**). That is, mechanical compression of the bacterial cell promotes persistent growth during bacteriophage predation.

**Figure 2.**
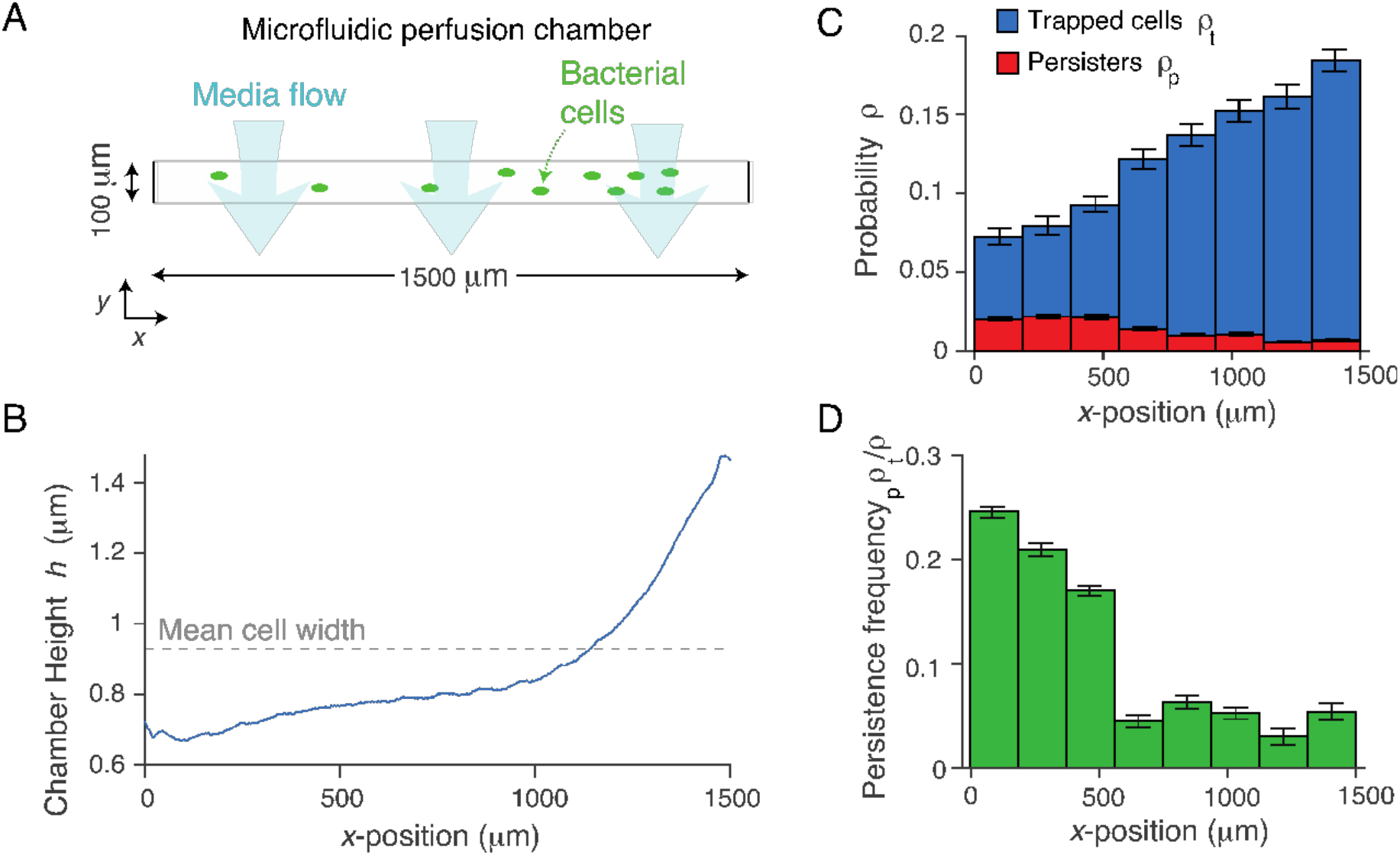
Persistent growth is inversely correlated with the degree of mechanical confinement. A) Schematic of microfluidic perfusion chamber. The chamber is rectangular with a large aspect ratio, and media perfuses parallel to the shorter (*y*) dimension. B) Height of a perfusion chamber versus position along the longer (*x*) dimension of the chamber. C) Mean spatial probability distribution of trapped cells (before bacteriophage perfusion) and persistent colonies (after bacteriophage perfusion) versus position in the perfusion chamber across six technical replicates. Error bars indicate ±1 s.d. D) Probability that a trapped cell would develop into a persistent microcolony versus position in the perfusion chamber. Error was propagated from data in (C).

### Rcs activation is required for compression-mediated persistence

We hypothesized that mechanical compression was inducing a transcriptional response that was protecting the bacteria from the bacteriophage, and moreover that the dynamics in colony size observed during persistence (**Fig. 1C,S1A**) reflected the dynamics of this transcriptional response. In a previous genome-wide screen, only a single gene, when overexpressed, provided resistance of *E. coli* to bacteriophage T7^27^. This gene, *rcsA*, encodes a transcription factor that, upon hetero-dimerization with a cognate transcription factor, RcsB, induces expression of an operon that synthesizes capsule, a polysaccharide expressed on the bacterial cell surface^28^. Furthermore, mutations that lead to constitutive overproduction of capsule result in bacteriophage resistance since capsule blocks the bacteriophage from binding its receptors in the outer membrane^29^. We therefore hypothesized that compression-mediated persistence was dependent on capsule synthesis (**Fig. 3A**). To test this, we measured the frequency of persistence in mutant bacteria that either lacked the ability to export capsule (Δ*wza*) or constitutively expressed capsule-synthesis enzymes (*rcsC137*)^30^. As hypothesized, capsule-less mutants were completely unable to grow persistently during bacteriophage predation, while constitutive capsule synthesis led to a dramatic increase in the frequency of persistent microcolonies (**Fig. 3B**).

**Figure 3.**
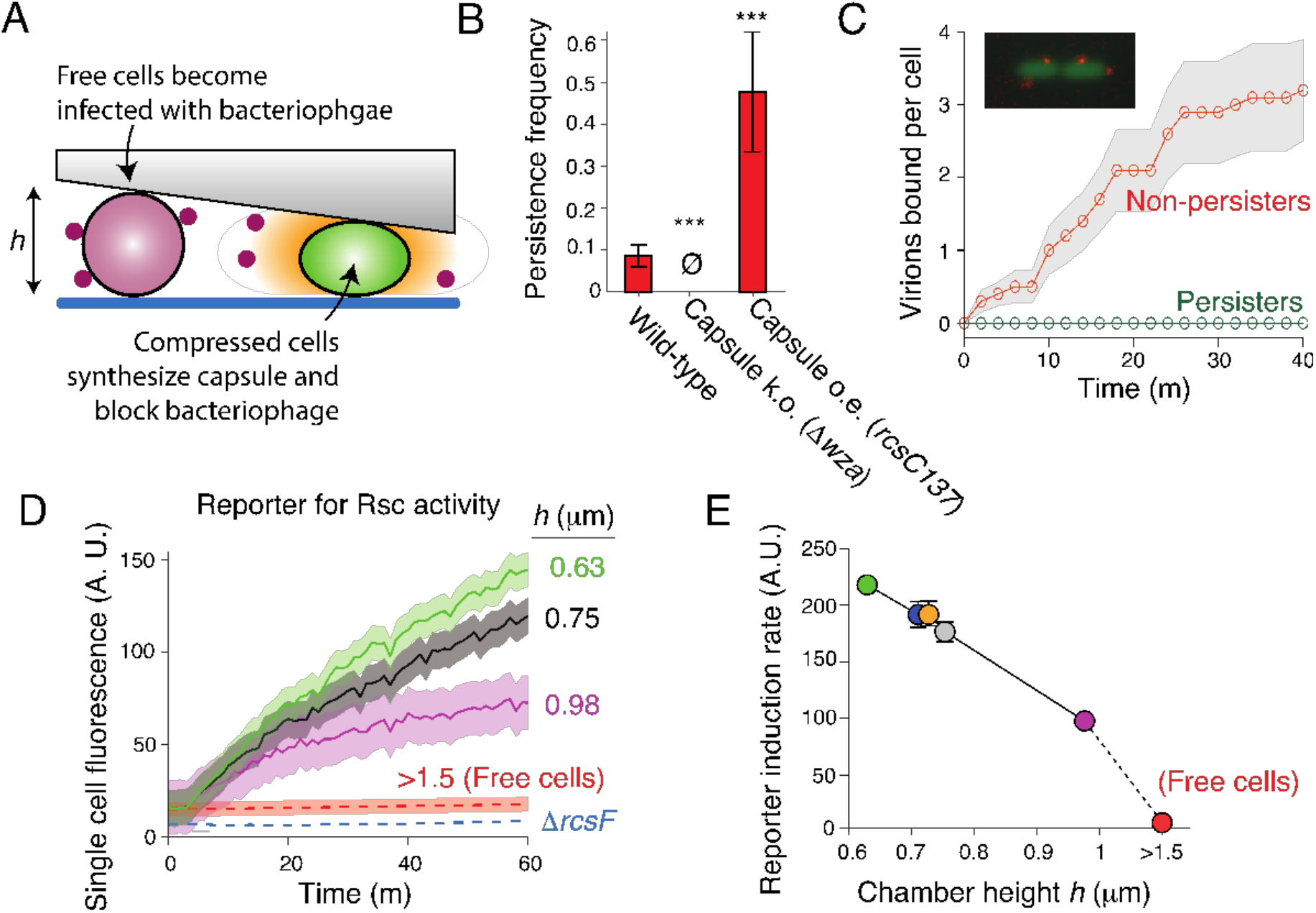
Mechanical compression induces Rcs activation. A) Model for bacteriophage persistence. B) Mean persistence frequency for wild-type *E. coli* and two isogenic mutants that either do not express capsule (Δ*wza*) or overexpress capsule (*rcsC137*). Error bars indicate ±1 s.e.m. across three technical replicates. 0: no persistent colonies observed. **: *p* < 0.001. **: *p* = 0.01. C) Population-averaged number of virions bound per cell versus time for cells that developed into persisters (non-lysing), and cells that did not, during bacteriophage perfusion. Confidence intervals indicate ±1 s.d. across *n* = 10 cells. (inset) Micrograph of two *E. coli* cells each with two bacteriophage T7 virions bound to them. D) Population-averaged single-cell fluorescence versus time and perfusion chamber height for *E. coli* cells harboring a plasmid-based transcriptional reporter for Rcs activity. Confidence intervals indicate ±1 s.d. across *n*=231,627,533,312,480,329 cells at each respective height. E) Population-averaged single-cell induction rate of reporter for Rcs activity versus perfusion chamber height, calculated from data shown in panel (E). Error bars indicate ±1 s.d. *n*=231,627,533,312,480,329 cells at each respective height.

To confirm that capsule synthesis prevented bacteriophage adsorption, we perfused *E. coli* with fluorescently labeled bacteriophage^15^ (**Fig. 1D**) and measured adsorption dynamics in persistent and non-persistent cells (**Fig. 3C**). We found that bacteriophage rapidly bound non-persistent cells (0.08 ± 0.06 virions cell^-1^ minute^-1^; **Fig. 3C,D**), but did not bind cells that developed into persistent microcolonies (**Fig. 3C,D**). Furthermore, bacteriophage adsorbed more slowly to non-persistent cells that constitutively synthesized capsule than to non-persistent wild-type cells (**Fig. S2**). Together, these data directly demonstrate the requirement of capsule synthesis for bacterial persistence to bacteriophage.

### Mechanical compression causes signaling through the Rcs pathway

Our results led us to a model in which mechanical compression causes bacteriophage persistence by inducing capsule synthesis (**Fig. 3A**). Expression of *rcsA* and its downstream regulon is induced by signaling through the Rcs phosphorelay pathway^28^. This complex pathway is triggered by cell-envelope stress, such as that caused by cationic antimicrobial compounds^31^, and ultimately induces up to ≈20 genes (including many not required for capsule synthesis), depending on the specific type of stress^31,32^.

To test whether mechanical compression activates Rcs signaling, we first constructed a plasmid-based reporter in which expression of super-folder GFP is controlled by the promoter of *rprA*, a gene whose transcription is directly induced by Rcs signaling^33^. We then measured single-cell reporter expression for cells compressed to varying degrees. We found that reporter expression was strongly negatively correlated with chamber height (**Fig. 3E,F**). Furthermore, reporter expression of cells trapped in the tallest area of the perfusion chamber was ≈50% of that of cells trapped in the shortest area of the cell, while reporter activity in bulk liquid culture was negligible (**Fig. 3D**), demonstrating that growth on surfaces alone (in addition to global mechanical compression of the cell) is sufficient to elicit Rcs signaling. Mechanical compression did not activate Rcs signaling in a mutant bacterium lacking RcsF, the protein that activates signaling in response to outer membrane stress (**Fig. 3E**), demonstrating that compression mediates capsule synthesis via a canonical mechanism.

### Multiple types of mechanical stimulation can elicit Rcs signaling and induce persistence

Surface-sensing and mechanical compression are two of many forms of mechanical perturbations that bacteria may sense in their environment. Thus, we next questioned whether other mechanical stimuli induce Rcs signaling and bacteriophage persistence. We first tested the effect of fluctuations in extracellular osmolarity, a fundamental environmental perturbation that causes variation in the intracellular turgor pressure within cells^34^. To do so, we subjected cells to an oscillatory osmotic shock^35^ (**Fig. 4A**) immediately before bacteriophage perfusion. This treatment caused modest increases in both the rate of Rcs activation and the frequency of persistence (**Fig. 4B,C**).

**Figure 4.**
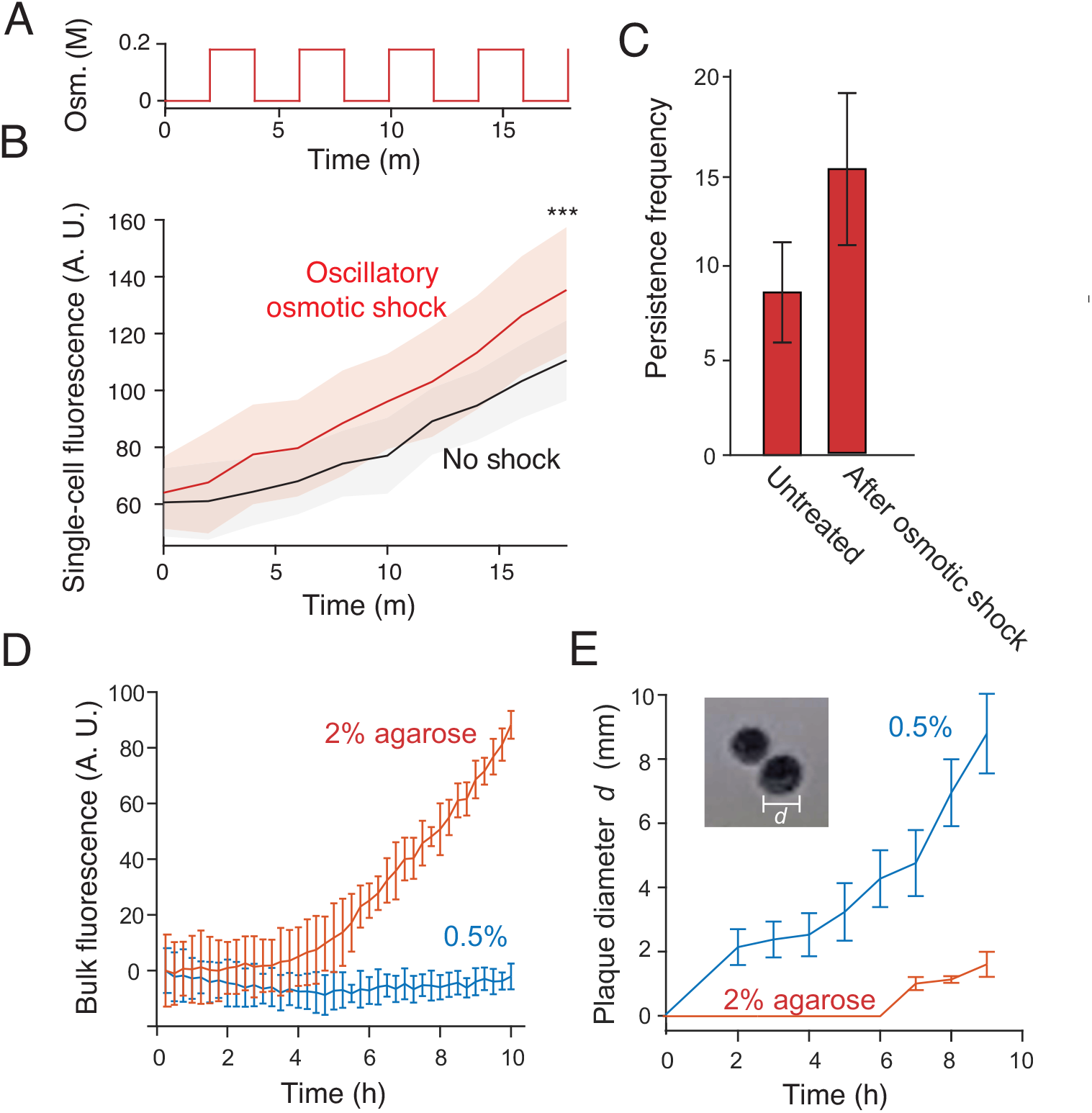
Several forms of mechanical stimulation cause persistence. A) Medium osmolarity versus time during a 200-mM oscillatory osmotic shock with 4-min period. B) Population-averaged single-cell fluorescence versus time for untreated (non-shocked) *E. coli* cells harboring a plasmid-based transcriptional reporter for Rcs activity and for the same strain undergoing an oscillatory osmotic shock as shown in (A). Confidence intervals indicate ±1 s.d. across *n*=35 and 15 cells for the two conditions, respectively. ***: *p* < 0.0001. C) Mean persistence frequency for untreated (non-shocked) *E. coli* and for the same strain after having undergone an oscillatory osmotic shock as shown in (A). Error bars indicate ±1 s.e.m. D) Mean bulk fluorescence versus time of *E. coli* cells harboring a plasmid-based transcriptional reporter for Rcs activity that are suspended in solid media containing different concentrations of agarose. Error bars indicate ±1 s.d. across *n*=3 technical replicates. E) Mean number of bacteriophage plaques per petri dish versus time when a single bacteriophage solution was diluted into suspensions of bacteria in solid media made with a range of agarose concentrations. Error bars indicate ±1 s.d. across 3 and 7 plaques for 2% and 0.5% agarose, respectively. (inset) Micrograph of two bacteriophage plaques. F) Mean plaque size versus time when a single bacteriophage solution was diluted into suspensions of bacteria in solid media containing 2% and 0.5% agarose. Error bars indicate ±1 s.d. across three technical replicates.

We next wondered whether persistence at the single-cell level could affect large-scale bacterial population dynamics. To do so, we used a bulk-culture method^36^ to assay the effects of mechanical stimulation on Rcs activation and bacterial proliferation during bacteriophage predation. First, we suspended cells in agarose of varying concentration and assayed Rcs activation in our reporter strain using a fluorescence plate reader. Similar to the case of mechanical compression (**Fig. 3F**), the rate of induction was much higher when cells were grown in stiff, 2% media than in soft, 0.5% media (**Fig. 4D,E**). Although we could not directly quantify single-cell persistence from this bulk-culture assay, we quantified the rate of bacteriophage proliferation by measuring the growth rate of bacteriophage plaques on these solid-media bacterial suspensions (**Fig. 4F**). As predicted, the rate of plaque growth was much slower on bacterial suspensions in stiff media than in soft media (**Fig. 4G**). Together with the increase in persistence in response to oscillatory osmotic shock (**Fig. 4B,C**), this result demonstrates that various types of mechanical stimulation, in addition to growth in confined environments, can induce persistence.

## Discussion

Here, we describe a new mechanism by which mechanically stimulated bacteria can grow persistently - and indefinitely - during lytic bacteriophage predation. They accomplish this by producing extracellular capsule in response to the mechanical stimulus, thereby slowing the dynamics of predation. In doing so, both the bacteria and the bacteriophage can proliferate simultaneously in the same environment.

We called this phenomenon “persistence” because it shares the essential attribute of antibiotic persistence^37^: that bacteria can survive the threat without a specific mechanism of resistance against it. Just as toxin/antitoxin-mediated antibiotic persistence^38^ is not a specific defense against antibiotics (but rather a generic mechanism for stochastic dormancy), capsule-mediated bacteriophage persistence is not a specific defense against bacteriophage, but a generic response to mechanical stimulation that may serve other purposes as well. It will be interesting to investigate whether mechanically stimulated capsule synthesis, in particular, has other functions. For example, the Rcs pathway promotes persistent *Salmonella* infections^46^, which could rely on the sensitivity on this pathway to mechanical stimulation in the gut. It is also possible that bacteriophage persistence is cross-protective against certain toxic compounds to which capsule is impermeable^39^.

In one sense, capsule-mediated persistence is similar to the protection that bacteria gain from bacteriophage by living in biofilms. In both cases, environmental stimuli induce the synthesis of an extracellular factor that inhibits bacteriophage replication and thus biofilms provide a form of persistence. However, the most remarkable feature of mechanically induced bacteriophage persistence, which distinguishes it from both biofilm-based bacteriophage protection and antibiotic persistence, is that it allows exponentially growing bacteria to survive bacteriophage. On one hand, this feature serves a clear function for non-resistant bacteria by providing niches (*e*.*g*., solid media or strongly fluctuating environments) where they can rapidly proliferate in spite of bacteriophage. On the other hand, this also allows bacteriophage to proliferate without extinguishing their host. Together, these characteristics of bacteriophage persistence promote stable coexistence of the two microbes.

Phylogenetically, the Rcs phosophorelay pathway is specific to enteric commensals and pathogens^46^, a group of bacteria that would have a clear need for the ability to transduce mechanosensation. Capsule, however, is much more widespread. It seems likely that bacteria from other ecological niches, for example those living in the soil, have co-opted other envelope stress response pathways for mechanosensation as well; this will be an interesting subject of future study for which the Rcs pathway will be paradigmatic.

The molecular mechanism by which mechanical stimulation activates the Rcs pathway is an open question. However, since compression-mediated capsule synthesis requires RcsF, it is likely that compression causes outer membrane damage that resembles the outer membrane stress caused by cationic antimicrobial compounds such as colistin. However, because Rcs activation is sensitive to very slight mechanical perturbations (*e*.*g*. growing on a surface, Fig. 3E,F) that do not inhibit rapid cell growth, this dependence may represent an adaptive form of mechanosensation mediated by forces in the outer membrane, rather than an emergency stress response. If so, this would add to a growing body of evidence that bacteria are exquisitely sensitive to their mechanical environment and execute sophisticated transcriptional programs upon mechanosensation^40^. For example, *Pseudomonas aeruginosa* executes a virulence program when it senses surfaces^41^; since in certain cases capsule is a virulence factor, our findings suggest that mechanical compression may also contribute to bacterial pathogenesis. Indeed, since bacteriophage are often present during bacterial pathogenesis^42^, protection from bacteriophage may be required for virulence in certain cases. On this note, our finding has major implications for the prospects of phage therapy: the ability of the therapeutic bacteriophage to overcome capsule must now be considered.

Bacteriophages are present in most natural environments where bacteria thrive^43^: in the gut, the soil, and virtually all bodies of water. In most of these environments, bacteria are directly subjected to mechanical forces either by other cells, by their environment, or via osmotic fluctuations. We propose that mechanically induced bacteriophage persistence is a previously unappreciated process that underlies the co-existence of bacteria and bacteriophage in many, if not most, of these environments.

## Materials and Methods

### Bacterial strains and culture conditions

All strains used in this study are listed in Table S1. All experiments were conducted with *E. coli* MG1655 or isogenic mutants. Bacteria were cultured in lysogeny broth (LB) with appropriate antibiotic selection at 37°C with shaking at 180 RPM, unless otherwise noted.

### Bacteriophage propagation and purification

Bacteriophage T7 was propagated using standard techniques. Autoclaved molten LB top agar (0.5 % agarose) was equilibrated to 65°C in a water bath. Serial 10X dilutions of bacteriophage T7 suspensions were made in bacteriophage buffer (10 mM TRIS pH 7.5, 10 mM MgSO_4_, 70 mM NaCl). 10µl of each bacteriophage dilution was added to 200µl of overnight *E. coli* MG1655 culture and incubated at room temperature for 20 minutes. 5mL of molten top agar was added to each culture and poured over a pre-warmed LB plate. Plates were incubated overnight at 37°C. The following day, 5ml of bacteriophage buffer was added to the top of the plate that had the highest number of individual plaques that could still be counted, and was incubated at room temperature for 2 hours. The phage buffer was then removed with a syringe and sterilized with through a 0.2 µm syringe filter.

### Strain and plasmid construction

All plasmids used in this study are listed in Table S2. Strain ER396 (MG1655, Δ*wza::Kan*) was generated by P1 transduction between ER373 (BW25113, Δ*wza::Kan*) and MG1655. Strain ER473 (MG1655, Δ*rcsF:Kan*) was generated by P1 transduction between ER367 (BW25113, Δ*rcsF::Kan*) and MG1655.

Plasmid pGM04 (genotype) was generated by Gibson assembly ^44^ of PCR amplicons of the backbone of pZS21^45^, the promoter and ribosomal biding site of rprA (60 base pairs upstream of the *rprA* start codon), and monomeric super-folder GFP (msfGFP). The *rprA* promoter was amplified from *E. coli* MG1655 and msfGFP was amplified from NO34^47^.

### Persistence assay

Prior to an experiment, 1 mL of LB was inoculated with 10 μL of overnight *E. coli* culture and incubated at 37°C with shaking for 2 hours. The cell inlet well of a CellASIC bacterial microfluidic plate (B04A) was loaded with 200 μL LB; two media perfusion wells were loaded with 200 μL of LB and LB containing a high-titer (≥10^7^ mL^-1^) of bacteriophage T7. The plate was then pre-warmed at 37°C in the microscope incubation chamber for 45 minutes, after which 2 μL of the exponentially growing bacterial culture was added to the loading well (1:100 dilution). The perfusion wells were then primed for 30 minutes in the microscope incubation chamber. The ONIX Microfluidic perfusion system was then used to load cells into the perfusion chamber at high pressure (10 psi) to drive cells into areas of the plate that had low chamber heights. Trapped cells were perfused with LB for defined periods of time, followed by indefinite perfusion of LB containing bacteriophage.

Bacterial cell growth and reporter fluorescence was recorded using a Nikon Ti2 inverted microscope and a Photometrics sCMOS camera. Fluorescence was excited using Aura Light Engine LED light source (Lumencore).

To subject cells to oscillatory osmotic shock prior to measuring persistence, LB containing 200 mM sorbitol was also loaded into a third media perfusion well on the microfluidic plate. After loading cells into the perfusion chamber, they were subjected to 10 cycles of a 200-mM oscillatory osmotic shock with 4-min period before LB containing bacteriophage was perfused indefinitely (with no osmotic shock).

### Perfusion chamber height determination

To determine perfusion chamber height before a persistence experiment, a fluorescent tracer dye (AlexaFluor 647 succinimidyl ester) was perfused into the chamber and a single stitched epifluorescence micrograph was taken of the entire chamber at 100X magnification. The intensity of the tracer dye as a function of *x*-position across a single slice (*y*-position) of the perfusion chamber was computationally measured from this image. To calibrate the intensity profile with the chamber height, upon loading cells into the chamber during the subsequent experiment, we found the largest *x*-position (corresponding to the tallest chamber height) at which cells became trapped and calibrated that height with the diameter of the cells, which we measured from the micrographs.

### Fluorescent labeling of bacteriophage

Covalent fluorescent labeling of the capsid of bacteriophage T7 was performed as described by Vidakovic *et al*.^15^. Briefly, 100 µl of a purified bacteriophage (10^11^ pfu mL^-1^) mixed with sodium bicarbonate (0.1 M final) were incubated with 0.1 mg AlexaFluor 647 succinimidyl ester for 1 hour at room temperature under continuous shaking and protected from light. The reaction mixture was dialyzed against PBS to separate bacteriophages from unbound dye for two days using 20,000 MWCO slide-a-lyzer cassettes. Fluorescently labelled bacteriophages were stored at 4°C.

Bacteriophage binding was counted manually.

### Computational image analysis

Tracking of the size of persistent microcolonies and quantification of single-cell fluorescence in the Rcs reporter was performed using custom MATLB scripts.

### Bulk culture measurements of Rcs reporter fluorescence and plaque formation

To measure induction of Rcs signaling in the reporter strain ER450 growing in solid media of varying stiffness, 200 μL molten LB media containing 0.5% or 2% agarose was equilibrated to 50°C and mixed with 10 μL of overnight bacterial culture in a 96-well plate. Fluorescence measurements were taken with a Tecan Spark Fluorescence plate reader at 15 minute intervals.

To measure plaque formation in agar of varying stiffness, LB agar plates were pre-warmed to 37°C and molten LB media containing 0.5% or 2% agarose was equilibrated to 50°C. 200 μL of overnight bacterial culture was mixed with 20 μL of bacteriophage T7, serially diluted to 1000 pfu/ml, and was incubated with the bacteria at room temperature for 20 minutes. 5 ml of molten agarose media was added to the culture tube and the mixture was poured over the pre-warmed agarose plates. Photos of the plaques were taken with a digital camera every hour for 9 hours.

## Acknowledgments

The authors would like to thank Kevin Rome and Andrew Darwin for technical assistance with molecular techniques, and Susan Gottesman and Thomas Silhavy for bacterial strains and plasmids. This research was supported by NIH grant 5R35GM143057-02 to E.R.R.

## Supporting Information for

**Fig. S1.**
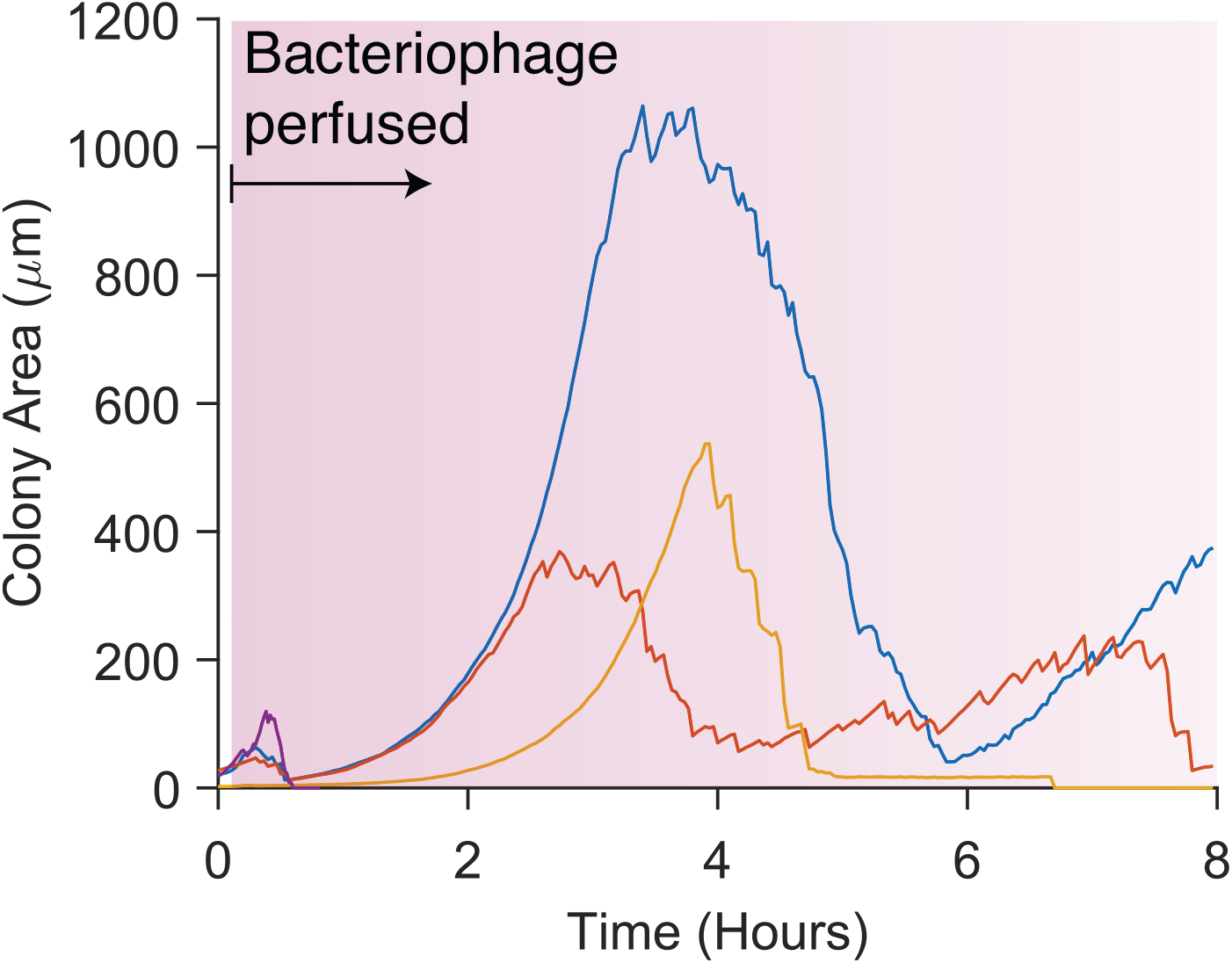
Persistence can occur after as little as 5 minutes of growth in the perfusion chamber. Colony area versus time for three *E. coli* microcolonies that grew under LB perfusion for 5 minutes before being perfused with LB containing bacteriophage T7 indefinitely.

**Fig. S2.**
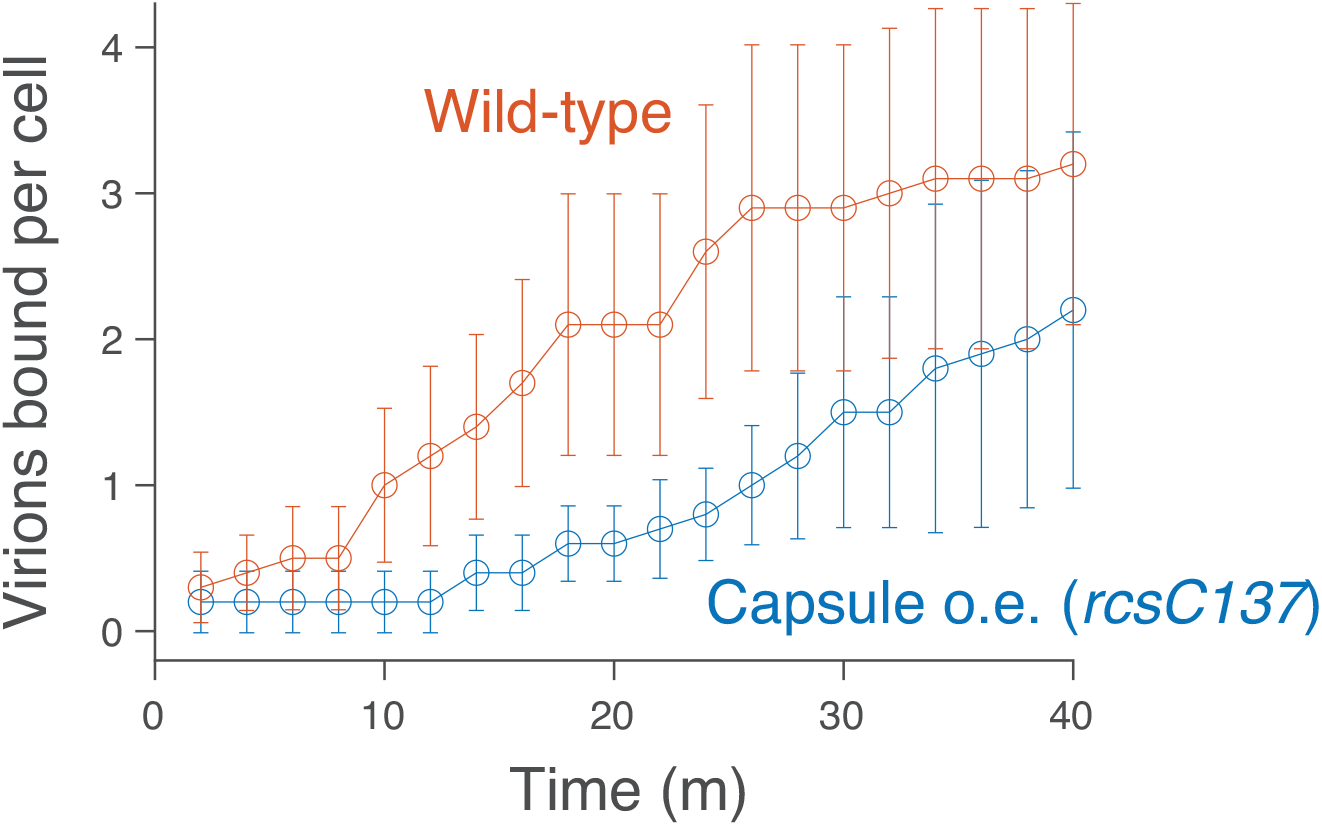
Capsule expression slows the rate of bacteriophage binding in non-persistent cells. Population-averaged number of virions bound per cell versus time for non-persistent wild-type cells and non-persistent mutant bacteria that constitutively overexpressed capsule (*rcsC137*). Confidence intervals indicate ±1 s.d. across *n* = 10 cell for each strain.

**Table S1.**
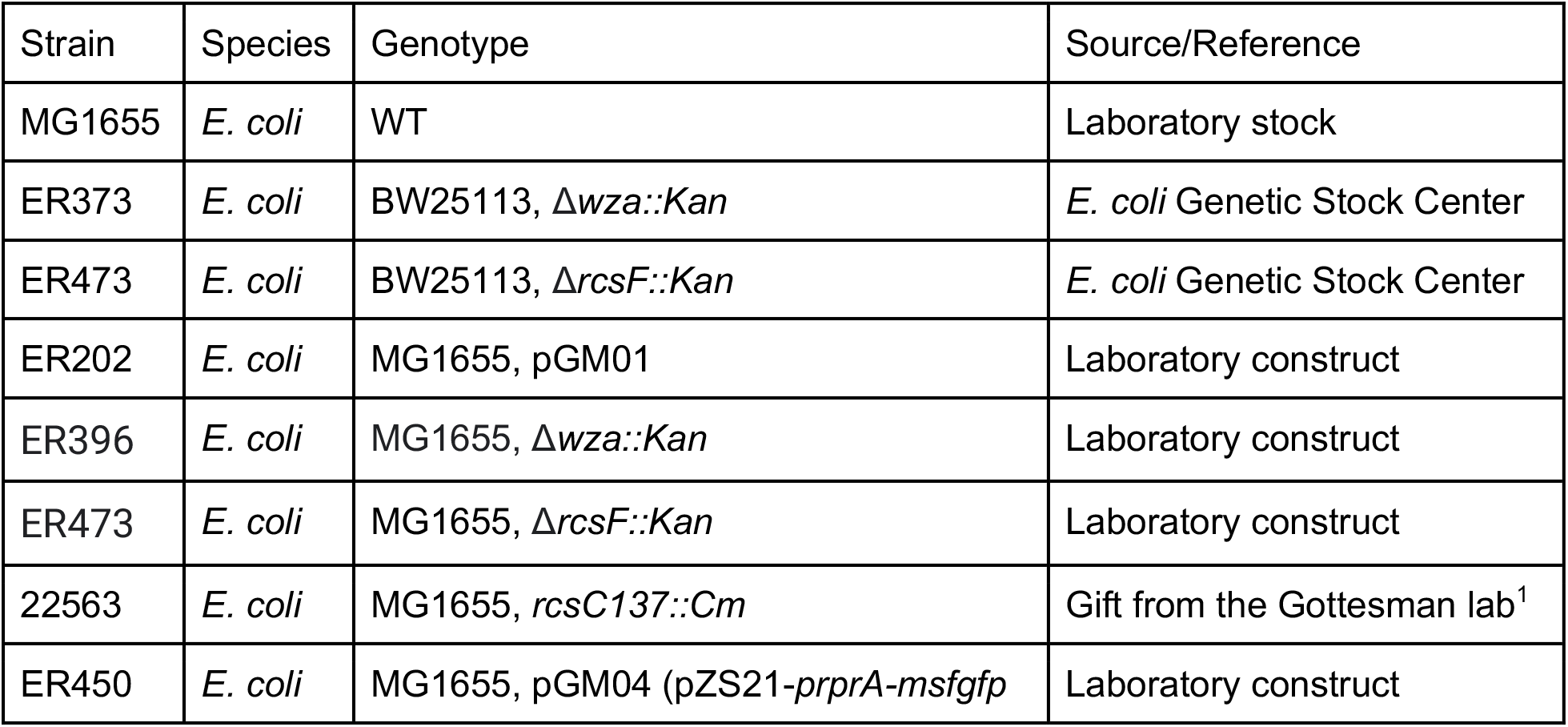
Bacterial strains used in this study.

**Table S2.**
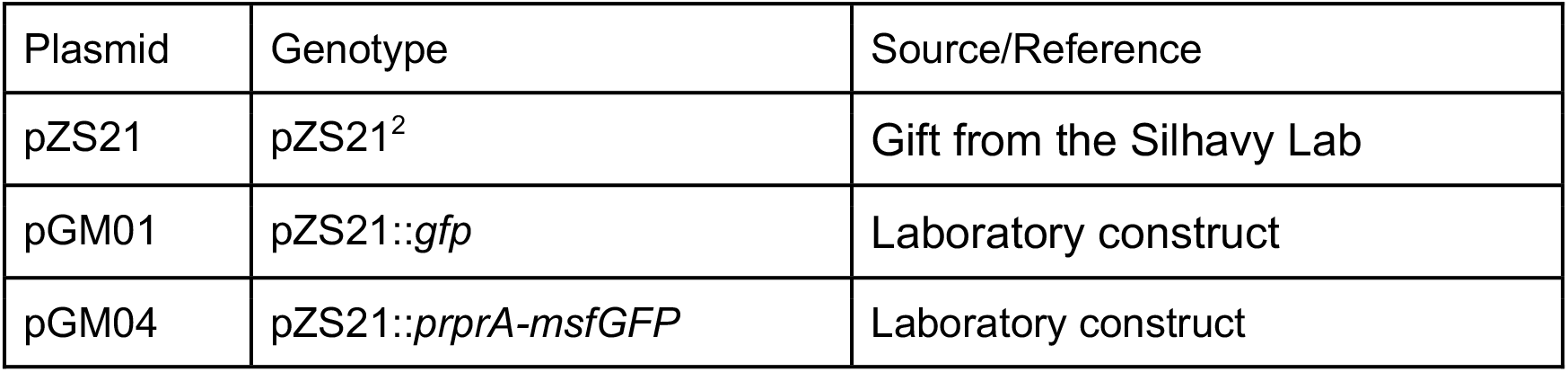
Plasmids used in this study

| Plasmid | Genotype | Source/Reference |
| --- | --- | --- |
| pZS21 | pZS21<sup>2</sup> | Gift from the Silhavy Lab |
| pGM01 | pZS21::*gfp* | Laboratory construct |
| pGM04 | pZS21::*prprA-msfGFP* | Laboratory construct |

**Movie S1 (separate file)**. Time-lapse micrograph (100X magnification) of *E. coli* cells expressing cytosolic GFP growing in a microfluidic perfusion chamber for 4 hours before being perfused with bacteriophage (not visible) for 11 hours.

**Movie S2 (separate file)**. Time-lapse micrograph (10X magnification) of *E. coli* cells expressing cytosolic GFP growing in a microfluidic perfusion chamber for 5 minutes before being perfused with bacteriophage (not visible) for 3 hours. Persistent microcolonies are enriched on the left side of the image where the height of the perfusion chamber is smallest.

